# Disentangling associations between complex traits and cell types with *seismic*

**DOI:** 10.1101/2024.05.04.592534

**Authors:** Qiliang Lai, Ruth Dannenfelser, Jean-Pierre Roussarie, Vicky Yao

**Author notes:** Correspondence to:* { }, { }.

## Abstract

Integrating single-cell RNA sequencing (scRNA-seq) with Genome-Wide Association Studies (GWAS) can help reveal GWAS-associated cell types, furthering our understanding of the cell-type-specific biological processes underlying complex traits and disease. However, current methods have technical limitations that hinder them from making systematic, scalable, interpretable disease-cell-type associations. In order to rapidly and accurately pinpoint associations, we develop a novel framework, *seismic*, which characterizes cell types using a new specificity score. We compare *seismic* with alternative methods across over 1,000 cell type characterizations at different granularities and 28 traits, demonstrating that *seismic* both corroborates findings and identifies trait-relevant cell groups which are not apparent through other methodologies. Furthermore, as part of the *seismic* framework, the specific genes driving cell type-trait associations can easily be accessed and analyzed, enabling further biological insights. The advantages of *seismic* are particularly salient in neurodegenerative diseases such as Parkinson’s and Alzheimer’s, where disease pathology has not only cell-specific manifestations, but also brain region-specific differences. Interestingly, a case study of Alzheimer’s disease reveals the importance of considering GWAS endpoints, as studies relying on clinical diagnoses consistently identify microglial associations, while GWAS with a tau biomarker endpoint reveals neuronal associations. In general, *seismic* is a computationally efficient, powerful, and interpretable approach for identifying associations between complex traits and cell type-specific expression.

## Introduction

Genome Wide Association Studies (GWAS) have shown exceptional promise for identifying genetic variants across populations that are key contributors to human diseases and a range of phenotypic traits. Simultaneously, the rise of large-scale single-cell RNA sequencing (scRNA-seq) datasets has revolutionized our ability to analyze gene expression profiles at the level of individual cell types and states. These advancements present a unique opportunity to integrate the population-level genetic associations revealed by GWAS with the molecular precision offered by scRNA-seq to pinpoint specific trait-associated cell types. Intuitively, a natural way to unify these datasets is to link trait-associated variants identified from GWAS to genes, which can then be subsequently analyzed for cell-type specificity using scRNA-seq.

Several computational methods have been developed to identify trait-associated cell types [1–6] by integrating reference scRNA-seq atlases with single nucleotide polymorphisms (SNPs) identified in GWAS. We focus on the major class of methods that use MAGMA [7] to account for the complexities of linkage disequilibrium. MAGMA itself can be adapted to predict cell type-trait relationships [3, 6], which we term S-MAGMA (see Methods) to avoid confusion with the linkage disequilibrium correction process. Broadly, these MAGMA-based methods have been successfully used to help elucidate disease-associated cell types [3, 6, 8–10], but several technical limitations remain (Table 1). One such limitation is the requirement of arbitrary thresholds, either for selecting the number of genes associated with a GWAS trait (e.g., top 1,000 trait-associated genes as in scDRS [1]) or to characterize a cell type (e.g., top 3,500 marker genes for S-MAGMA [3] as used in [10]), or even for defining the specific score quantile for assessing cell-type association enrichment (e.g., upper 5% quantile as in scDRS). Biologically, these numbers would naturally vary case-by-case, and most methods recommend trying several. Methods also often fail to account for gene expression variability, focusing instead more simply on mean expression within a cell type, rendering them more susceptible to noise. Furthermore, as the size of single cell datasets continue to grow, scalability becomes a major concern, necessitating a method that can handle large numbers of cells and cell types. Finally, a critical gap in previous methods is that though they may identify several significant cell type-trait associations, they only output statistical significance scores without further gene-level interpretation. Although some methods attempt to resolve the issue using global correlation [1] or modularity analysis [4], they fail to quantify each gene’s contribution to the observed significance association of an interested cell type, limiting the actionable insights to be derived from these analyses.

**Table 1.** Comparison of *seismic* with other MAGMA-based cell-type association methods.

|  | *seismic* | scDRS | FUMA | S-MAGMA |
| --- | --- | --- | --- | --- |
| No gene thresholding based on trait association | ✓ |  | ✓ | ✓ |
| No gene thresholding based on cell-type expression | ✓ | ✓ | ✓ |  |
| Accounts for gene expression variability | ✓ | ✓ |  |  |
| Scalable runtime | ✓ |  | ✓ | ✓ |
| Tool includes results visualizations | ✓ | ✓ | ✓ |  |
| Identifies cell-type level influential genes | ✓ |  |  |  |

Here, we present *seismic*, a framework that enables robust and efficient discovery of cell type-trait associations and provides the first method to simultaneously identify the specific genes and biological processes driving each association. Notably, *seismic* eliminates the need to select arbitrary thresholds to characterize trait or cell-type association through the use of a cell-type-specificity scoring method that accounts for background gene expression variability. We apply *seismic* and existing MAGMA-based cell-type association methods on both simulated and real data to demonstrate that *seismic* is well-calibrated, efficient, and powerful.

Through a deep exploration of neurological disease-associated brain cell types, we find that cell type definition from input scRNA-seq data is an important, underappreciated, factor that influences downstream findings. Specifically, previous analyses [6, 8, 11, 12] have typically used broad characterizations of cell types such as “telencephalon projecting excitatory neurons” and “frontal cortex neurons,” which do not consider additional region or tissue granularity, possibly obscuring valuable biological insights. For example, in Alzheimer’s disease, a population of neurons in the entorhinal cortex are most vulnerable, while neurons in even neighboring brain regions such as the dentate gyrus and CA2/CA3 in the hippocampus are not [13]. We find that finer granularities for cell type characterization translates to higher resolution associations that better reflect true biological associations. Notably, *seismic* is more consistent than other methods at identifying disease-associated cell types across these different cell type characterizations. Finally, we demonstrate the importance of considering different GWAS endpoints to reveal disease mechanisms, reporting, to our knowledge, the first identification of a neuronal association with an Alzheimer’s disease biomarker (tau level in cerebrospinal fluid). Together, our results expand current notions of best practices for cell type-trait association analyses and provide a methodological toolkit to take fuller advantage of both scRNA-seq and GWAS data to unravel the intricate interplay between tissue/cell type and complex traits.

## Results

### Identifying cell type-trait associations using *seismic*

Many cell type-trait association methods consider the same inputs—variant-trait information from GWAS resolved to gene-trait using MAGMA [7] and single cell expression data—to find statistically significant associations between cell types and traits (Figure 1A). Here, we introduce a novel integration framework, Single-cell Expression Investigation for Study of Molecular Interactions and Connections (*seismic*), that overcomes the limitations of previous methods to provide a threshold-free, fast, and interpretable method for combining single cell expression data with gene-trait relationships (Figure 1B).

**Figure 1.**
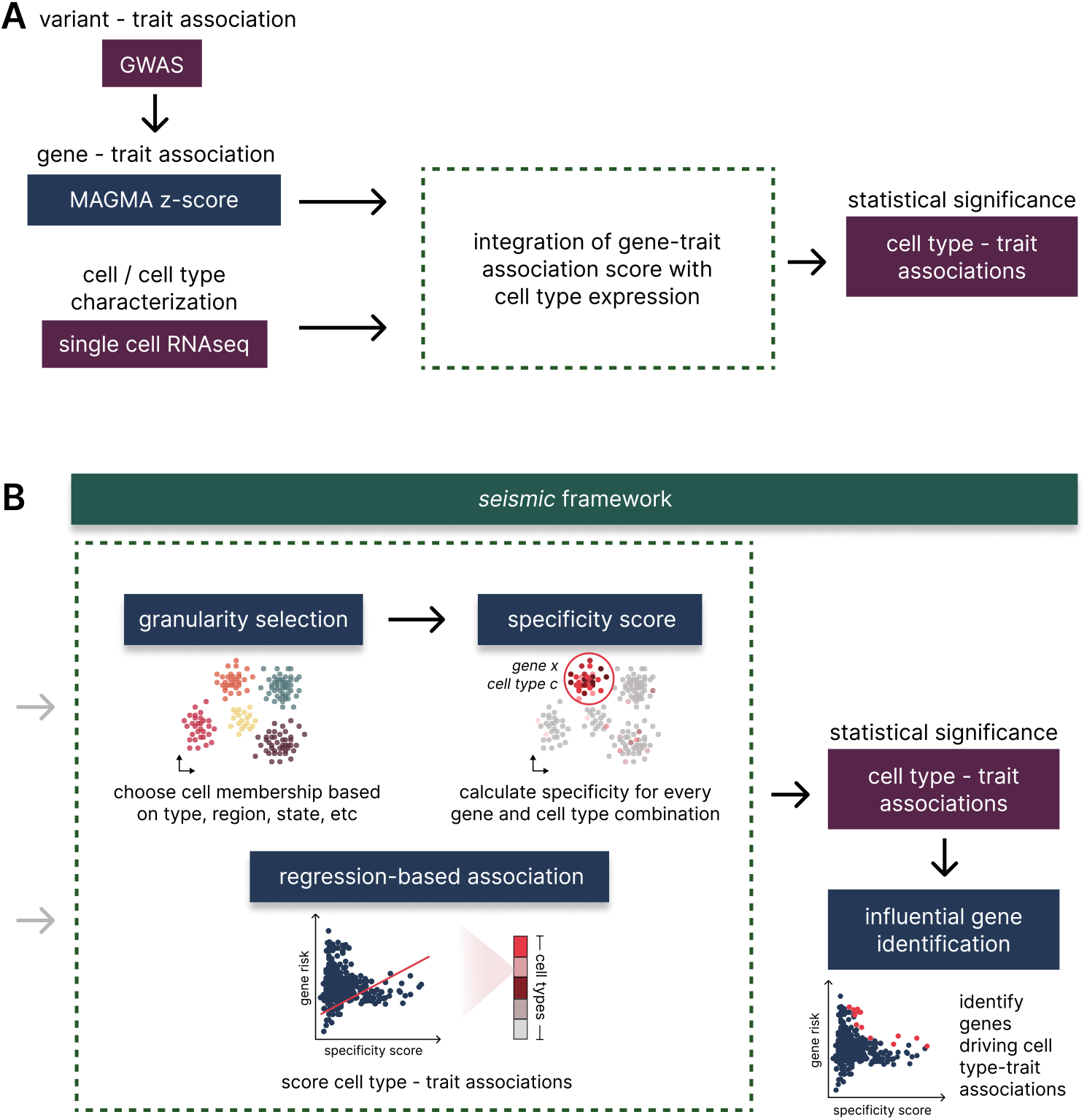
General framework for identifying cell type-trait associations and overview of *seismic*. **(A)** Current cell type-trait association methods integrate single cell expression data with gene-trait associations from MAGMA to prioritize cell type-trait associations by statistical significance. Each method has a different procedure for how these inputs are combined to produce the final set of identified associations. **(B)** The *seismic* framework expansion of this general workflow. *seismic* allows for a flexible set of cell type labels or granularities, ranging from broader cell classes to specifically defined cell types, before calculating a novel gene specificity score for each of the corresponding cell characterizations. Within the selected granularity, *seismic* calculates the statistical significance of a cell type-trait association using a regression-based framework with the gene specificity scores and MAGMA z-scores (Methods). Unlike existing methods, individual gene contributions to the cell type-trait association can be quantified via influential gene analysis and can pinpoint the genes and underlying biological processes that drive significant associations.

At the core of *seismic* is a cell-type-specificity score (Methods) which calculates the specificity and consistency of expression for each gene in a cell type relative to all other cell types. For a collection of cell types in a scRNA-seq dataset, *seismic* then uses a regression model to test for significant associations between the gene specificity scores and MAGMA gene z-scores. We find that *seismic*’s cell type-specificity scores enable it to easily adapt to broad and precise definitions of cell type (granularities), allowing for analyses using both more inclusive cell type labels, such as “immune cell” or “neuron,” as well as highly specific cell type characterizations incorporating region, cell type, and even cell state, such as “CA1 pyramidal cell” or “reactive microglia.” For these significantly associated cell types, the *seismic* framework also applies influential observation analysis to the respective regression model, enabling what we term ‘influential gene analysis,’ which is, to our knowledge, the first method to systematically rank and identify genes driving purported cell type-trait associations.

### Systematic benchmarking for false discoveries and runtime

To assess how well *seismic* and three of the most commonly used cell type-trait identification methods (scDRS [1], FUMA [2], and S-MAGMA [3]) are calibrated to false positives, we perform a systematic simulation to detect the frequency of type I errors. We first randomly select 10 sets of MAGMA trait z-scores from GWAS (Supplementary Table 1) and subsample 10 expression datasets, each containing 10,000 cells from the Tabula Muris FACS scRNA-seq data [14]. For each subsampled expression dataset, we randomly select 100 cells as a cell type of interest (Methods). Next, across 10,000 runs, we randomize the gene labels in the expression data and compare the p-values reported by each method for the association between the randomly assigned target cell type and trait. We find that all methods generally control type I error, with FUMA being markedly conservative, potentially limiting its detection power. *seismic* is on average conservative and has stable performance. In contrast, using the analytically transformed p-values from scDRS, we see slightly inflated p-values at extreme quantiles, and S-MAGMA can also at times report inflated p-values (Figure 2A). The *seismic* and scDRS implementations enable examination of the effect of randomization of MAGMA trait z-scores, and we observe the same trends, where *seismic* has generally well-calibrated p-values, and scDRS’ has slight inflation at higher quantiles (Supplementary Figure 1).

**Figure 2.**
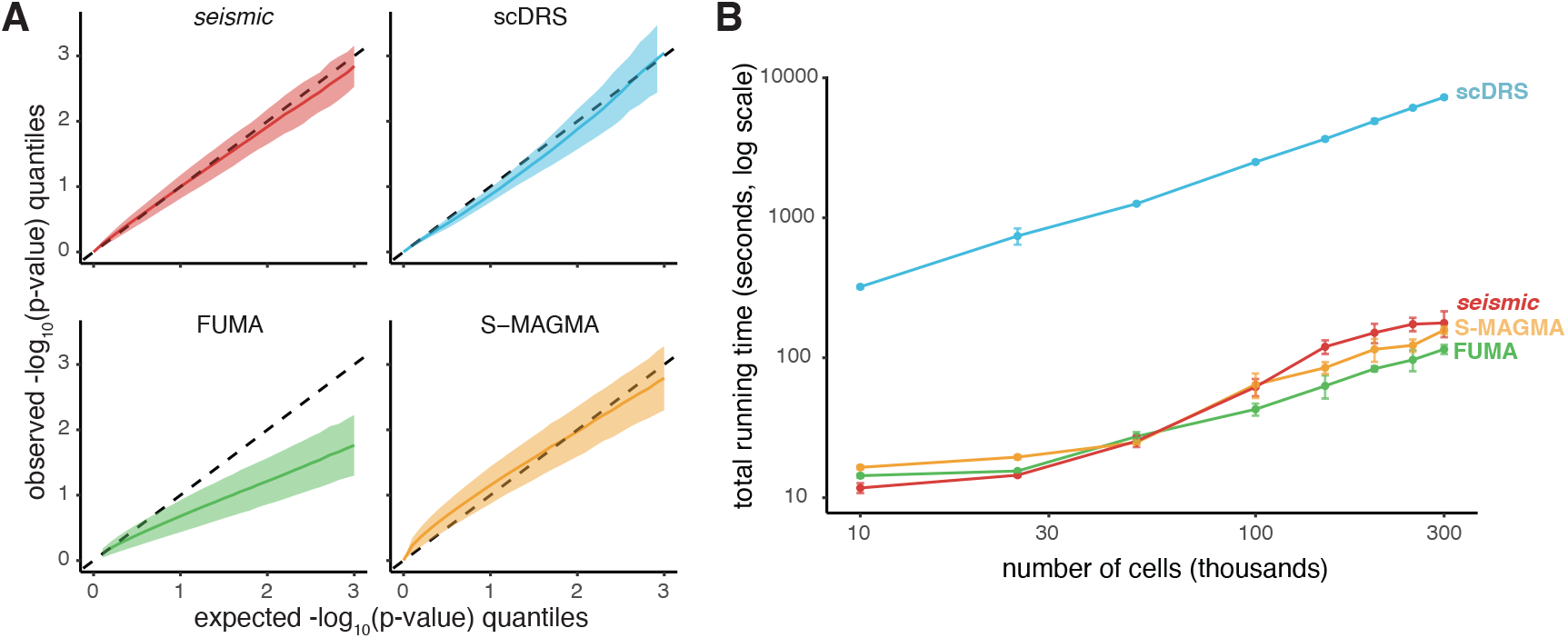
Systematic benchmarking results for *seismic* and three other commonly used methods for cell type-trait association detection. **(A)** Assessment of type I error calibration for *seismic*, scDRS [1], FUMA [2], and S-MAGMA [7]. Results are based on random cell type assignments and expression datasets across 10,000 simulation runs with 10 randomly selected traits and 10 subsets of 10,000 cells (Methods). Quantile-quantile plots show the comparison of expected − log_10_(p-value) quantiles in comparison with observed − log_10_(p-value) quantiles, so the dashed line represents perfectly calibrated p-values, and observations falling below the dotted line indicate that p-value estimates are more conservative. Ribbons denote the standard deviation of the − log_10_(p-value) quantile. **(B)** Total running time for each method to assess the trait association for all cell types in the dataset in seconds (shown in log scale) as the number of cells increases. *seismic* demonstrates comparable runtimes to S-MAGMA and FUMA and scales well as the number of cells increase, unlike scDRS, likely due to the Monte-Carlo-based subsampling procedure it uses for statistical estimation. Error bars denote the standard deviation of the runtime across samples.

As single cell technologies continue to improve and we move closer towards atlas-scale datasets, computational methods need to be able to scale well with the number of cells. To this end, we also benchmark the four methods for their runtime as the number of cells in the input single cell expression dataset increases (Figure 2B, Supplementary Table 3). *seismic*, FUMA, and S-MAGMA scale comparatively well, handling hundreds of thousands of cells with runtimes in the scale of minutes, whereas scDRS is over 30 fold slower, taking hours to run. We speculate that the dramatic difference in speed between scDRS and all other methods is likely due to its reliance on Monte Carlo-based subsampling for empirical statistics, whereas all other methods directly assess associations at the cell type level and do not rely on simulations for significance assessment. Together, these benchmarking results highlight *seismic*’s ability to appropriately account for both type I errors while scaling well to the size of the single cell expression input.

### Methodological comparisons across traits and cell types

To examine *seismic*’s ability to capture known cell type-trait associations across a broad range of GWAS traits, we assemble 27 studies spanning neurological diseases and disorders, immune-related conditions, and a variety of other traits, including demographic, cardiovascular, and metabolic endpoints (Supplementary Table 1). We test for cell-type associations using the expression values and annotations from the Tabula Muris FACS dataset [14], which includes nearly 45,000 cells, covering 130 cell type characterizations across 17 tissues (Supplementary Table 2). In total, we find 653 pairs of cell type-trait associations that pass the significance threshold of FDR *≤* 0.05 (Supplementary Table 4). Notably, *seismic* identifies associations linking leukocytes with immune diseases, neurons with neuropsychiatric diseases, smooth muscle cells with cardiovascular diseases, pancreatic cells with type 2 diabetes, and hepatocytes with metabolic traits (Figure 3A), recapitulating known biological cell type-trait associations. Moreover, *seismic* is robust against variations in gene window size (average correlation between pairs > 0.98, Supplementary Figure 2).

**Figure 3.**
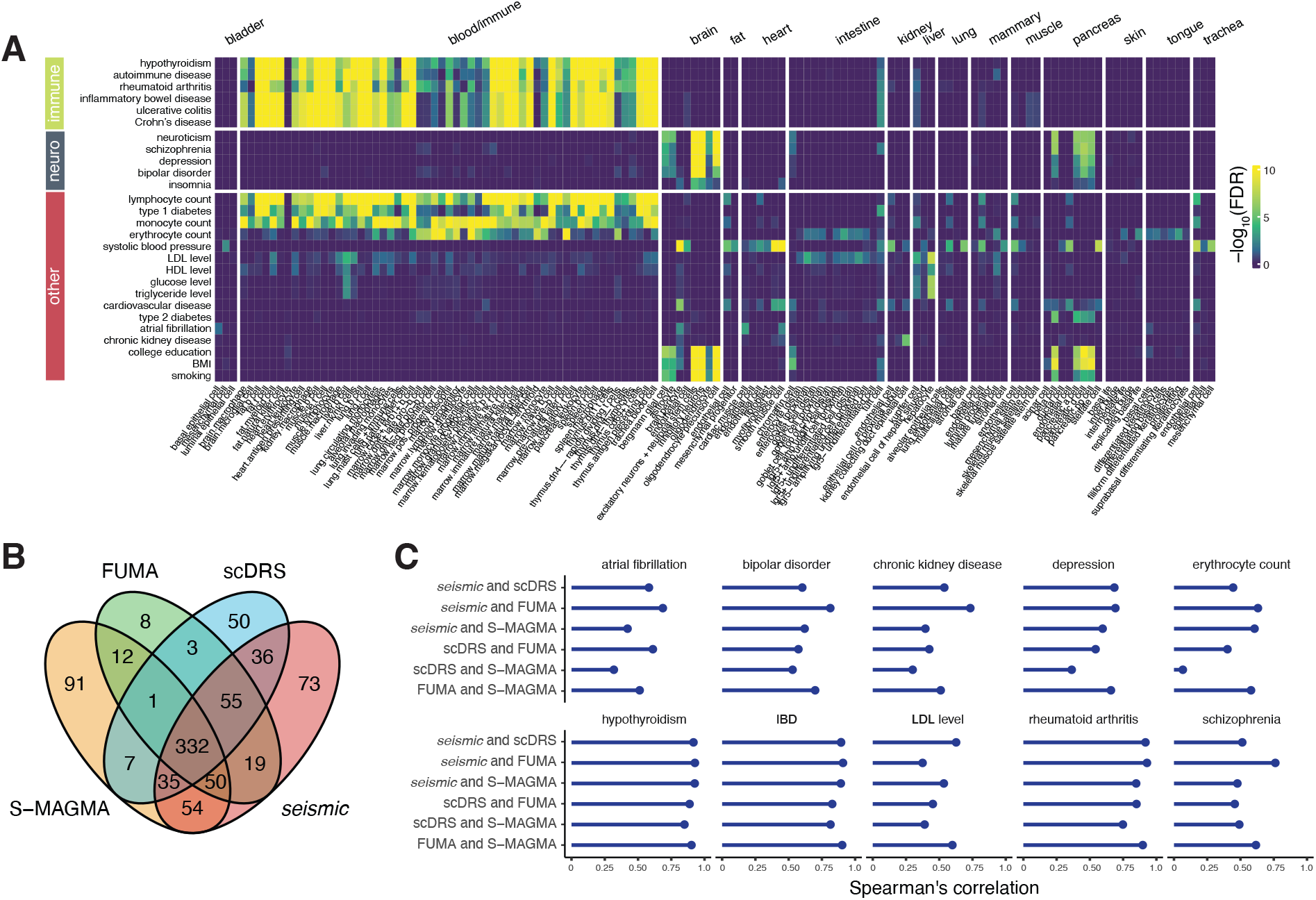
*seismic* finds biologically relevant cell type-trait associations across a variety of GWAS studies. **(A)** We apply *seismic* on 27 GWAS studies covering neurological, immune, and various other disease or demographic endpoints with gene expression from the Tabula Muris FACS dataset [14] to identify cell type-trait associations covering 15 tissue types. While there is no gold standard for these associations, we find many expected cell type-trait relationships. **(B)** Venn diagram comparing the significantly associated cell type-trait relationships between methods (based on a 0.05 FDR threshold). *seismic* identifies many overlapping cell type-trait relationships with existing methods, while finding 73 additional associations missed by other methods. **(C)** Pairwise Spearman’s correlations for each of the methods across a selected set of 12 diverse traits (other trait correlations are in Supplementary Figure 10). In general, *seismic* has the highest correlation with other methods.

We then apply S-MAGMA, FUMA, and scDRS to the same dataset and GWAS traits, checking for consistency with *seismic*’s results and any differences in associations (Supplementary Table 4, Supplementary Figures 3-8). Notably, 89% of associations identified by *seismic* are also detected by at least one of these frameworks (Figure 3B, Supplementary Figures 6-8), where *seismic* captures most of FUMA’s reported associations (95%), followed by scDRS (88%), then S-MAGMA (68%). This high degree of overlap highlights *seismic*’s robustness and alignment with established methods. The high correlation is further illustrated in a detailed between-method comparison, which reveals that *seismic* consistently achieves the highest trait-wise concordance among the methods, as measured by Spearman’s correlation. Specifically, for 26 of the 27 traits examined, *seismic* and one other framework achieves the highest concordance (Figure 3C, Supplementary Figure 10). Notably, *seismic* shows the highest average between-method Spearman’s correlation across all traits (0.69), compared with 0.61 for scDRS, 0.66 for FUMA, and 0.60 for S-MAGMA. For the 332 common association pairs found by all frameworks, *seismic* exhibits the most significant false discovery rates (FDR) in over 81% of these pairs (270 out of the 332 pairs), demonstrating its power (Supplementary Table 4).

Beyond these common findings, *seismic* also identifies additional association patterns that may better capture the underlying biology. For erythrocyte count, while scDRS ranks several cells from the intestine as most relevant, *seismic* and S-MAGMA identify several hematopoietic lineage cell types in marrow to be most associated, more accurately reflecting the developmental process of red blood cells. *seismic* also observes broad associations between neuropsychiatric diseases and various pancreatic islet cell types, which is especially noticeable in depression. These somewhat outlandish associations are also recapitulated by the other methods, and interestingly, previous studies have found potential associations between pancreatic and neuropsychiatric diseases [15–18], which has led to increased interest in a potential pancreas-brain axis. In total, *seismic* only misses 1 cell type-association pair identified by all other methods (Figure 3B, Supplementary Table 4), the fewest compared to other methods (50, 35, and 55 pairs for scDRS, FUMA, and S-MAGMA, respectively). The undetected association, between microglia and autoimmune disease, is close to the multiple hypothesis threshold (with FDR=0.066, Supplementary Table 4).

To explore generalizability to other large scRNA-seq datasets, we also apply *seismic* to the Tabula Muris (TM) droplet dataset [14] (a dataset obtained by droplet-based single-cell sequencing rather than profiling individually sorted cells as in TM FACS) and the Tabula Sapiens (TS) human scRNA-seq dataset [19]. Comparing cell types that overlap between TM FACS and these 2 additional scRNA-seq datasets, one using a different technology, and the other using human cells, we find consistent cell type-trait associations (Supplementary Figures 9, 11, 12, mean Spearman’s correlation of 0.79 between TM FACS and TM droplet, and 0.67 between TM FACS and TS; we note that neither TM droplet nor TS contain brain tissue, and the mean Spearman’s correlation is 0.84 and 0.75 respectively if neuropsychiatric traits are excluded from the comparison). The high concordance of *seismic* with other methods is also consistent across datasets (Supplementary Figures 10, 13, 14). Such consistency underscores *seismic*’s robustness in identifying trait-associated cell types across datasets of larger size, varying coverage, as well as *seismic*’s adaptability to different species.

### *seismic* associations are consistent across tissue-cell type granularities

Having examined *seismic*’s consistency in identifying a wide-variety of trait-associated cell types, we turn our attention to evaluate the accuracy of *seismic’s* ability to distinguish known vulnerable neuron types for a well-characterized neurological disease, Parkinson’s disease (PD). PD pathophysiology is well-established, with dopaminergic neurons residing in the substantia nigra pars compacta (SNc) and ventral tegmental area (VTA) characterized as being particularly vulnerable to degeneration [22]. Using a large mouse brain dataset [21] encompassing up to 231 distinct cell types from 9 regions of the adult mouse brain, in conjunction with a recent PD GWAS study [20] with over 480,000 participants, we test whether *seismic*, scDRS, FUMA, and S-MAGMA can recover known PD associations (Figure 4, Supplementary Table 5). Additionally, the rich brain region and cell type annotations in [21] provide a unique opportunity to test how changes in cell type granularity affect the reported cell type-trait annotations. We examine 5 different granularities of cell types, ranging from 14 broad subclass labels to 231 highly-specific cell annotations (brain region + fine cluster). *seismic* is the only method to significantly prioritize PD-relevant dopaminergic neurons across all cell type granularities (Figure 4). FUMA and scDRS also rank relevant cell types in some granularities highly, but mostly fail to reach statistical significance after multiple hypothesis correction. Notably, S-MAGMA completely misses these vulnerable cell types. We note also that most previous cell type-trait association analyses that use datasets such as [21] typically perform their analyses at a broader cell type level (usually what we have termed the ‘brain region + class’ granularity). Though using finer resolution annotations increases the number of multiple hypotheses compared, we demonstrate that it may be a worthwhile trade-off, as it can lead to more precise biological insights.

**Figure 4.**
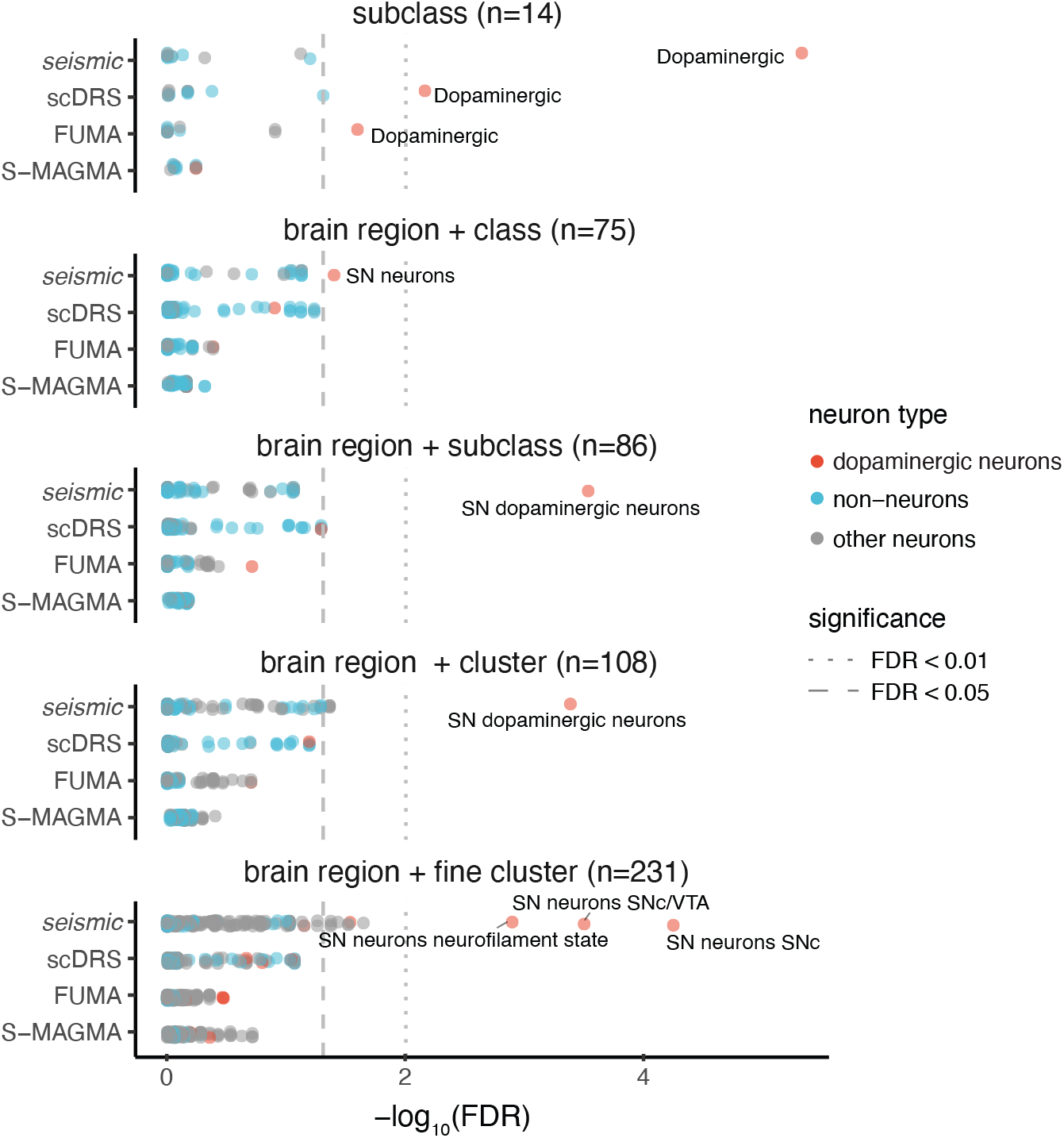
*seismic* finds Parkinson’s disease relevant cell types across granularities. A Parkinson’s disease GWAS study [20] analyzed using cell type labels at different resolutions from Saunders et al. [21]. Cells are partitioned into increasingly more specific cell types, the broadest of which is subclass labels (top), to brain region + fine clusters (bottom). *seismic* is the only method that correctly identifies Parkinson’s disease relevant neurons, dopaminergic (DA) neurons primarily localized to the substantia nigra (SN), across all cell type granularities. Any cell types with FDR < 0.01 are labeled, except for the subclass and brain region + class granularities, where cell types with FDR < 0.05 are labeled.

### Case study: applying the *seismic* framework to study Alzheimer’s disease

The choice of an endpoint for cell-type trait association can allow for the dissection of cell contribution to various endophenotypes of disease. This is particularly true for diseases with multicellular pathogenesis like Alzheimer’s disease (AD), where genetics studies have mapped clinical and alternative traits, making it also an ideal test case for demonstrating the power of *seismic*. Furthermore, while selective neuronal vulnerability and pathological legion formation have been thoroughly described in AD [23–26], the precise molecular mechanisms driving neurodegeneration leading to cognitive decline remain poorly understood. Formally, AD is characterized by two pathological hallmarks, extracellular amyloid plaques composed of A*β* peptide and intracellular neurofibrillary tangles (NFTs) formed by aggregated tau protein. NFTs appear according to a stereotypical spatial pattern, first emerging in the layer II of entorhinal cortex (EC), later appearing in deeper layers of EC and CA1 in the hippocampus, before subsequently spreading to other neocortical and subcortical regions. Progression of NFTs is accompanied by neurodegeneration in the affected area. In spite of the strong correlation between clinical symptoms of the disease and neuronal processes (NFT formation, neurodegeneration, synapse loss), many GWAS studies have primarily identified associations with immune cells such as microglia [27–29]. This leads to questions of whether microglia are the primary drivers of the disease or merely responsible for the clinical symptoms of the disease. If the latter, one would expect to find cell-type associations for GWAS with non-clinical, pathology-based endpoints, which could open new research avenues for understanding pathogenic mechanisms.

We use *seismic* to test whether GWAS for different AD-related endpoints might yield divergent cell-type associations. We first use an AD GWAS that includes a large cohort of >63,000 patients diagnosed via clinical observations (clinical GWAS) [30]. This large study is representative of the AD GWAS typically used in cell type-trait association studies. We also explore *seismic* results for a GWAS for an alternative AD endophenotype comprised of around 3,100 patient samples of cerebrospinal fluid (CSF) tau levels [31], which serves as a biomarker of AD progression (tau GWAS) [32]. Though the tau GWAS has a much smaller patient cohort, we hypothesized that it may deliver clues for pathological mechanisms that have remained elusive with clinical GWAS. Applying *seismic* on these two AD studies along with the expression data from Saunders et al [21], as with previous methods, we identify microglial cells from various brain regions as the most associated with clinical GWAS (Figure 5A), demonstrating the pervasive neuroinflammation patterns underlying clinical symptoms in AD patients. Interestingly, cell-type associations with the tau GWAS primarily captured specific neuronal signal corresponding to the neurons most vulnerable to tau accumulation in AD (Figure 5B), including deep (“HC neurons entorhinal cortex”) and superficial (“HC neurons medial entorhinal cortex 1”) layers of the entorhinal cortex, as well as CA1 pyramidal cells.

**Figure 5.**
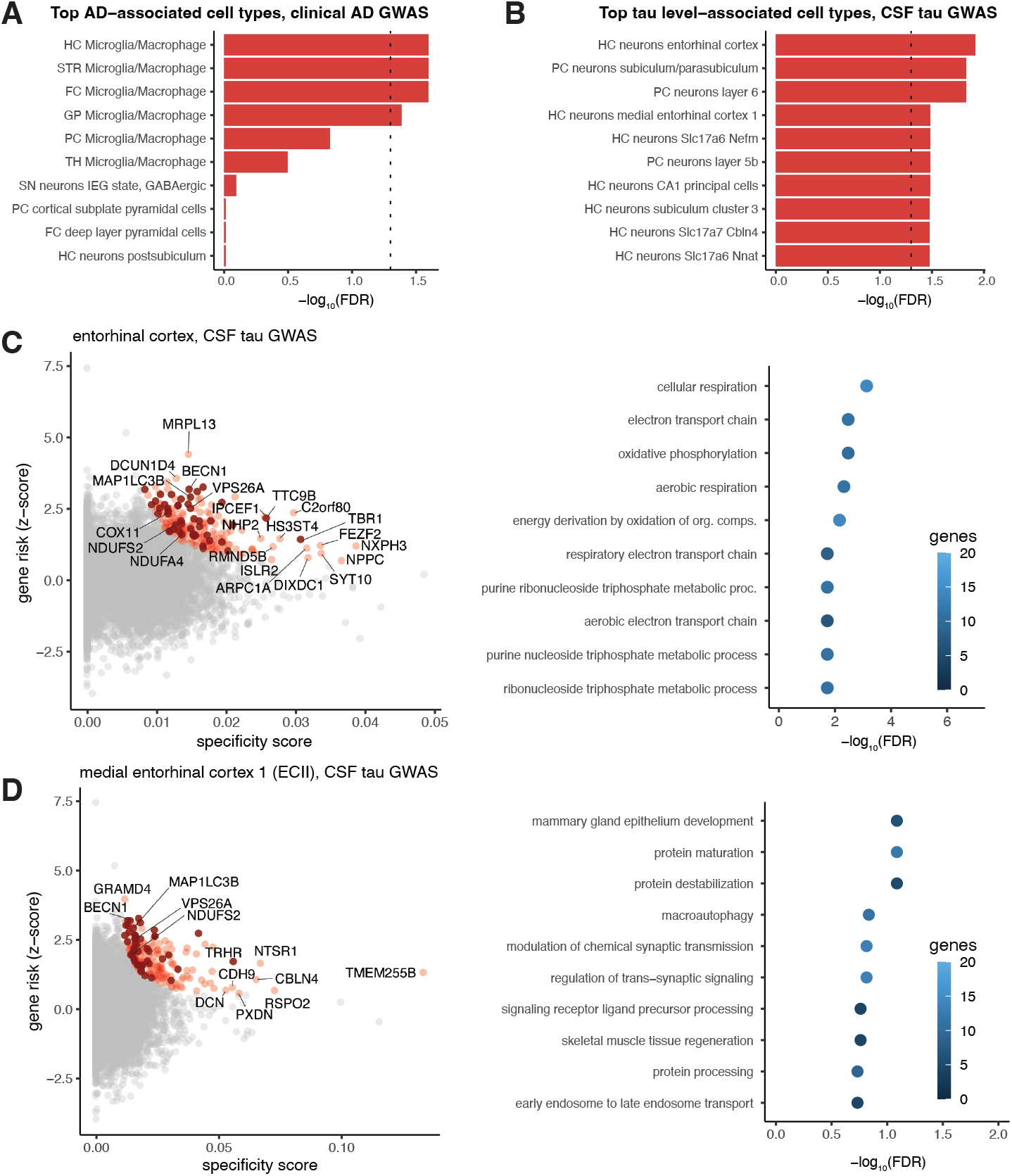
*seismic* enables the study of precise Alzheimer’s disease (AD) pathology. **(A)** *seismic* applied to AD GWAS study with a clinical endpoint reveals microglia cell associations (additional abbreviations: ENT, Entopeduncular Nucleus). **(B)** Swapping the AD GWAS using a clinical endpoint with one that measures an AD biomarker (CSF tau levels) reveals neuronal associations between the entorhinal cortex (EC) and other neurons in the hippocampus (HC) and posterior cortex (PC) (additional abbreviations and specifications: STR, striatum; “HC neurons entorhinal cortex,” EC excitatory neurons positive for Nxph3; “HC neurons medial entorhinal cortex 1,” EC layer II excitatory neurons positive for Cbln1). **(C)** Influential gene analysis for the genes driving the association of neurons froom deep layers of the entorhinal cortex (HC neurons entorhinal cortex) and the tau GWAS. Genes colored in red are influential, and dark red indicates influential genes driving the associations for both these and ECII neurons; several top influential genes are labeled. GO terms enriched for these genes are shown in the enrichment plot to the right. **(D)** Influential gene analysis for the genes associated with neurons in the medial entorhinal cortex (HC neurons medial entorhinal cortex 1, which we refer to as ECII) and the tau GWAS. As in C, influential genes are colored red, and influential genes shared between deep layers of the EC and ECII are darkened. The GO plot shows enrichment for the corresponding influential ECII genes.

With *seismic*’s influential gene analysis, we more closely inspect the genes corresponding to increased risk driving the top cell type-trait association. For hippocampal microglia, we identify 88 genes as positively influential for clinical AD diagnosis, and find these to be enriched for immune-related GO processes (Supplementary Figure 15, Supplementary Table 6). Some of these genes are expected, like SPI1, MS4A6A and TREM2, which are microglia-specific and known to have significantly reported associated SNPs with clinical AD. There are also several other interesting genes identified by *seismic*, such as LAPTM5, an amyloid plaque responsive gene [33], and phagocytosis regulator VAV1 [34]. Perhaps less expected are the influential genes found for entorhinal cortex neurons (218 genes for neurons from deep layers of EC, 199 genes for neurons from entorhinal cortex layer II) and the tau GWAS, 46 of which are found to be influential in both neuronal populations (Figure 5C, D, Supplementary Table 6).

In spite of the reasonably sized overlap of influential genes between neurons in different layers of the entorhinal cortex associated with the tau GWAS, we find distinct enriched pathways. Genes driving associations with CSF tau levels in deep layers of EC have generally more significant enrichment, primarily in metabolic pathways (e.g., cellular respiration, electron transport chain), while those driving associations with tau levels in layer II neurons are more enriched in proteostasis pathways (e.g., protein destabilization) (Figure 5C, D). Metabolic [35] and proteostatic [36, 37] contributions to vulnerability of EC are among the very pathways previously suggested to underlie EC vulnerability, and several identified influential genes, such as VPS26A, have connections with EC vulnerability in both cell types. The fact that *seismic* does not find any association between microglia and CSF tau further suggests that microglia might not be the main drivers of tau pathology, but rather the drivers of the clinical manifestations of AD. Meanwhile, the observation that using the tau GWAS enables *seismic* to not only identify neuronal associations, but also genes and pathways with great mechanistic and therapeutic potential, argues for the value of more targeted endophenotype GWAS—albeit smaller—for complex diseases.

## Discussion

Atlas-scale single cell RNAseq datasets have been generated with the promise of many exciting future applications. One way in which we can realize this potential is by combining them with quantitative genetic studies, helping to disentangle the tissue- and cell-specificity of complex traits and diseases. So far, several existing tools integrating scRNA-seq with GWAS studies to uncover cell type-trait associations have emerged [1–3], but these tools have several drawbacks (Table 1) that *seismic* addresses, including removing the need for thresholds, accounting for gene expression variability, and being scalable with the number of cells in an expression dataset. Importantly, *seismic* proposes influential gene analysis as a means to derive deeper biological insights for cell type-trait analyses.

Through our application of *seismic*, we also wanted to highlight the importance of cell type granularity selection and choice of GWAS endpoints, both of which seem to have a large impact on the final results, but, to our knowledge, have not been fully considered in previous analyses. Here, we observed that *seismic* can find results that are largely robust to changes in cell type granularity (Figure 4), while changes in granularity seemed to more strongly affect the cell type-trait associations detected by other methods both in the statistical power of detecting true associations and in the prioritization of specific cell types. We thus strongly recommend running any method at several cell type granularities to assess result robustness and using the finer cell type definitions to reveal more mechanistic interactions. Furthermore, as evidenced by the AD case study, it may be fruitful to include more targeted GWAS endpoints when studying complex disease. One of the interesting future directions we envision for *seismic* is an extension to automatically test for cell-type associations at different cell type granularities simultaneously. Such a feature would further complement the expanding cell reference atlases and facilitate faster identification of important cell-trait signal.

The application of *seismic* on the two AD-related GWAS studies (Figure 5) touches on a long mystery in the AD community, where large clinical GWAS consistently only identify microglial associations in spite of neuronal disease hallmarks and a strong selective neuronal vulnerability pattern not reflective of the relative regional homogeneity of microglial cells. Here, we show, to our knowledge, for the first time a neuronal association reflecting those most pathologically vulnerable to AD with an AD-related GWAS. This case study further highlights the exciting potential of *seismic* to find disease-relevant signal in small cohorts in a way that was previously impossible. In addition to homing in on a particular endophenotype, this case study also may reflect how GWAS using pathological or biomarker endpoints can intuitively avoid several pitfalls of clinical AD GWAS: inclusion of individuals with significant silent pathology in control groups [38] or mixed pathologies in the diseased group [39], as well as symptom gravity modulated by factors that have little to do with pathogenic mechanisms [40–42]. Tau levels in the CSF comes partly from tau secreted from neurons or from dying neurons, and should be influenced by processes taking place in neurons with significant tau accumulation and/or degenerating neurons. Association of this trait with neurons most vulnerable to NFT formation out of all brain neuron types profiled by Saunders et al. [21] confirms the power of the *seismic* framework, as well as the existence of neuronal signal in GWAS for alternative AD phenotypes which can yield mechanistic insight. The discovery that genes involved in metabolic and proteostatic processes underlies the association between EC neurons and CSF tau is exciting for a number of reasons. First, EC has been suggested to present notably high metabolic activity [35, 43]. Neurons from different layers of EC, in particular in the layer V, display a unique and energetically taxing phenotype of persistent firing activity [44], key for memory formation/consolidation process. On the other hand, neurons from layer II of the EC have a complex axonal arborization that might display high levels of structural plasticity, and thus might rely on high levels of protein turnover [45]. Aging-associated defects of proteostasis [46] would hit such neurons particularly hard [36]. Importantly, whether for metabolism or proteostasis, the influential gene analysis leads to experimentally testable hypothesis for the link between EC neuronal processes and vulnerability to tau accumulation.

We recognize that our method, as with previous cell type-trait association methods, has some fundamental limitations. Ultimately, statistical significance suggests a strong association, but not necessarily causality between cell types and traits. Even though in our study, *seismic* has demonstrated impressive performance in capturing trait-associated cell types that accurately reflect the corresponding biological knowledge, it may still miss some associations. For example, in the Tabula Muris FACS analysis (Figure 3), *seismic* missed 23 associations that are identified by at least two other methods. We do note that a closer inspection of these associations reveal that several may be spurious (e.g., Crohn’s disease, inflammatory bowel disease, and ulcerative colitis associations with limb muscle skeletal muscle satellite cell, all results in Supplementary Table 4), suggesting that not all of these 23 associations are necessarily false negatives. Another limitation is the usage of a scRNA-seq mouse brain atlas [21] in our PD and AD analyses, as opposed to a human scRNA-seq brain expression atlas. Unfortunately, current limitations in data quality and scale hinder us from using human brain-level data right now [47]. TS does not include brain tissue, but we do find that *seismic* identifies associations that are consistent between the mouse TM FACS dataset with the human TS dataset in overlapping cell types. We also note that as found in previous studies [1], the mouse datasets are often cleaner. As atlas-scale human data continues to improve, we expect the signal we can detect with *seismic* to also improve.

In conclusion, we have developed a new methodological toolkit to take better advantage of scRNA-seq and GWAS data to model the interplay between tissue, cell type and complex traits. We make all code for *seismic* and the accompanying analyses available through GitHub (https://github.com/ylaboratory/seismic-analysis) and installable as an R package, seismicGWAS (https://github.com/ylaboratory/seismic).

## Methods

### The *seismic* framework

To identify cell type-trait associations, *seismic* takes 2 inputs: (1) MAGMA [7] z-scores processed from GWAS summary statistics for a given trait; and (2) a scRNA-seq dataset that covers cell types of interest (Figure 1A). More details regarding how we processed GWAS and scRNA-seq datasets can be found below (see ‘Data preprocessing’). Many scRNA-seq datasets have inherent hierarchical structure (e.g., cells can be grouped by tissue of origin and also further divided by cell subclass or cell state). In other words, cell types can be categorized at different levels of *granularity*, for example adding resolution to a traditional cell type characterization by also considering its tissue sub-region. In most applications, we recommend choosing finer granularities, which typically translates to higher resolution results.

One of the motivating assumptions of *seismic* is that the genes that best characterize a cell type are not necessarily the genes that have highest expression in the cell type, but instead the genes that are *most specific* to that cell type. Optimally, these cell type specific genes would have consistently higher expression in all cells within the cell type and much lower or even no expression in other cells. They key insight of *seismic* lies in how to translate this optimal criterion into a continuous specificity score that can be calculated from a given scRNA-seq dataset and collection of cell types.

#### The *seismic* specificity score

The optimal criterion can be broken down into two sub-criteria: (1) consistently higher expression in a cell type of interest in comparison to other cells; and (2) expression in all cells within the cell type. These two sub-criteria are naturally related, but capture different aspects of cell-type-specificity. The first sub-criteria captures the variability and magnitude of expression, while the second focuses on the proportion of cells in a cell type where the gene is expressed (in a binary sense, ignoring the magnitude of expression). Here, we describe how continuous scores representing each of these sub-criteria are calculated.

Let *E* be an *N* × *M* matrix representing scRNA-seq expression data with *N* genes and *M* cells. With *j* ∈ {1, …, *M*} representing the index for each cell in *E* and *c* ∈ {1, …, *C*} representing the index for each of *C* cell types at the selected granularity, we represent cell type set membership as a labeled set of cells *L*^(*c*)^ = {*j*|*j* is labeled as cell type *c*}. For a given cell type *c*, we define *E*^(*c*)^ as the sub-matrix of *E* with column indices provided by *L*^(*c*)^. *E*^(*c*′)^ is the sub-matrix of *E* for the set complement of *L*^(*c*)^, which represents the expression of all cells not labeled as *c*.

We first seek to estimate the probability a gene has consistently higher expression in a cell type of interest. In other words, we want to estimate a score 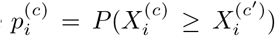, where 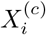 is a random variable representing the expression level of gene *i* in cell type *c*. We see that:

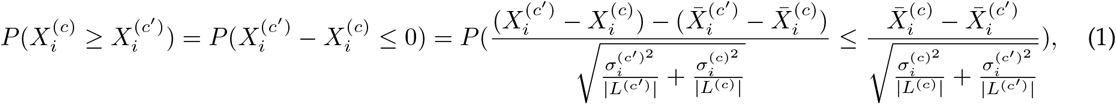

where 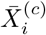 and 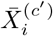 are the sample mean expression for gene *i* in cell type *c* and all other cells, respectively, 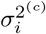 and 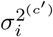 are the corresponding sample variances, and |*L*^(*c*)^| and |*L*^(*c*′)^| are the number of cells in cell type *c* and all other cells, respectively. Thus, if there are sufficiently large enough number of cells in cell type *c* (and otherwise), we know that based on the Central Limit Theorem, we can estimate 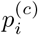 as:

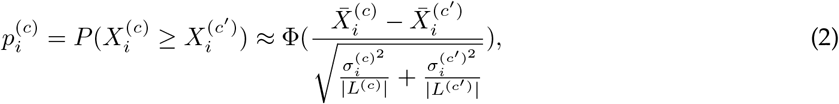

where Φ is the cumulative distribution function (CDF) of the standard normal distribution. We note that this formulation carries similarities with the derivation for the test statistic for a two sample z-test; however, we are primarily interested in calculating a good estimate for 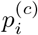 and not in calculating p-values that would correspond to a null hypothesis for no difference in expression. In this current formulation, 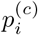 is bounded from [0, 1], where higher scores reflect higher expression in cell type *c* for gene *i*, even after considering the variability of expression.

Next, we consider how to estimate the probability of expression across cells in a cell type of interest if we consider expression as a binary variable. We see quickly that this is akin to estimating the ratio or proportion of cells in a cell type where we see non-zero gene expression. Thus,

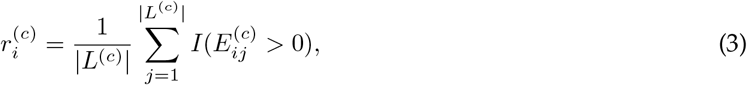

where *I* is the indicator function. As with 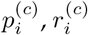 is also bounded from [0, 1], but here, higher scores reflect that gene *i* is expressed in a higher proportion of cells in cell type *c*.

Finally, we define the *seismic* specificity score as:

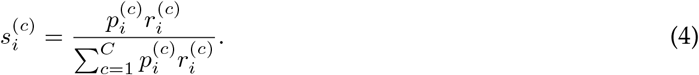

Here, we see that 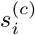 is still bounded between [0, 1], but now, 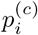 and 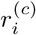 are also rescaled across all cell types to further highlight the specificity. A score close to 1 can be achieved if gene *i* both has consistently higher expression in cell type *c* compared with other cells even after considering expression variability and is also expressed in all cells within the cell type. As part of the *seismic* framework, we calculate the specificity score 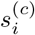 for all genes *i* and cell types *c* in the dataset.

#### Quantifying cell type-trait associations

We assume that if a cell type is highly associated with a trait, then the cell-specificity of its gene expression should have explanatory power for gene risk. In other words, if we observe that as the specificity of a gene to a particular cell type increases, there is correspondingly increased association with a disease, that would suggest a strong cell type-trait association. This can be formulated as a linear model:

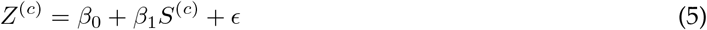

Here, *Z*^(*c*)^ denotes the vector of gene z-scores for cell type *c* given by MAGMA gene analysis, and *S*^(*c*)^ is the vector of *seismic* specificity scores for cell type *c*. Only genes that both have a z-score from MAGMA and are captured in the processed scRNA-seq datasets are considered.

After fitting the linear model, we can test the null hypothesis of *β*_1_ = 0 against the one-sided alternative hypothesis *β*_1_ *>* 0, resulting in a p-value for each cell type *c*. To correct for multiple hypothesis testing, we calculate and report Benjamini-Hochberg false discovery rates (FDRs). The one-sided test here tests for a positive linear relationship between cell-type-specificity and gene risk; we note that it is also possible that the relationship can be non-linear, which could potentially be addressed in the future using kernel methods. However, the advantage of this formulation is that we can statistically quantify the association in a directly interpretable way and enables influential gene analysis (below).

#### Influential gene analysis

For a significant cell type-trait association pair, some genes with may be particularly “influential” for the linear model. Here, we are using the statistical definition of an influential observation, specifically that removing the observation (i.e., gene) would have a strong effect on the model. Here, we calculate the difference in betas (DFBETAS) statistic [48], which is a scaled measure of how much the model parameters will change when removing a single observation. Specifically, for each gene and *β*_1_:

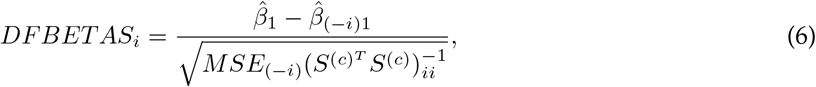

Where 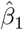 is the regression coefficient estimated with *all* genes, 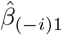 is the new regression coefficient calculated with gene *i* removed, *MSE*_(−*i*)_ is the mean squared error of the updated linear model without gene *i*, and 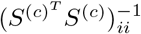 is the *i*^*th*^ diagonal element of the 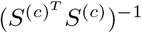 matrix calculated using *all* genes. As recommended by Belsley et al., we use the size-adjusted threshold for selecting top influential genes as 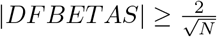, where *N* is the number of genes modeled [48]. For gene set enrichment analysis of the resultant influential genes, we use clusterProfiler [49].

### Existing MAGMA-based cell type-trait association methods

For scDRS [1] runs, we use the default parameters (1000 genes) with 1000 Monte Carlo (MC) samples and the top 5% quantile of trait scores across cells within a cell type as the test statistic for a given cell type. Because scDRS requires disease gene sets and an expression dataset as input, our processed data (‘Data preprocessing’) required some additional processing. Disease gene sets were generated from MAGMA z-scores using the provided *munge-gs* command. For cross-species analyses, scDRS deals with cross-species gene matches by their gene names; as such, we annotated the gene name entries of the MAGMA z-score files according to biomaRt mappings between entrez IDs and gene symbols [50]. If a gene is mapped to multiple z-scores, we collapse by calculating the mean gene z-score. Rare cell types (those with fewer than 20 cells) were removed from expression datasets to enable consistent comparisons across methods. scDRS recommends the use of empirical P-values for assessment of statistical significance; however, the resolution is limited by the number of MC simulation runs, especially in a multiple hypothesis test setting. Thus, we use the alternate recommendation of transforming disease-cell type association MC z-scores to their analytical p-value, followed by FDR calculations.

FUMA [2] provides a website for GWAS preprocessing and cell type-trait association analysis, but it is challenging to systematically analyze new datasets, especially for the null simulations and runtime analyses. We thus processed scRNA-seq datasets into the input format required by MAGMA (with expression values represented as log(CPM+1) as in FUMA) and used FUMA’s recommended parameters and commands to obtain p-values for each trait and cell type using the MAGMA software.

MAGMA-specific gene set analysis (S-MAGMA [3]) takes as input the non-log-transformed CPM expression. For S-MAGMA comparisons, as in Bryois et al [6], we used rescaled gene expression (where each gene’s expression value is divided by the total expression of that gene across all cell types), and the top 10% of genes were used for downstream testing.

### Null simulation

Null simulations were performed using random subsamples of the Tabula Muris (TM) FACS dataset [14] and randomly selected GWAS endpoints. More specifically, we randomly selected 10 GWAS endpoints from the 27 GWAS traits examined in Figure 3 (Supplementary Table 1), then randomly subsampled 10 subsets of 10,000 cells from TM FACS to use as input expression data for each method, where 100 cells were randomly selected as the target cell type. In our first null simulation setup, for each of 10,000 runs, the gene indices for the expression matrix were randomized. This random shuffle essentially breaks any biologically-relevant signal between the single cell expression and the gene z-scores. Keeping the MAGMA z-score fixed, the p-value for the random cell type and trait association was calculated for each method and compared with the expected p-values.

We also performed the more direct null simulation using shuffled z-scores on *seismic* and scDRS. (FUMA and S-MAGMA were excluded from this comparison because they require complete MAGMA gene analysis files (as opposed to more easily manipulated z-score files as in *seismic* and scDRS).) In this setup, for each of the 10,000 runs, we randomized the MAGMA z-score vector used as input to *seismic* and scDRS together with the same randomly selected target cell types from before (without further randomization of the expression matrix) and examined the p-values reported for cell type-trait association.

### Runtime analysis

We sampled a range of 10,000 to 300,000 cells from the Tabula Sapiens [19] (TS) data set to compare the end-to-end runtime for identifying associated cell types from a scRNA-seq dataset for 5 random traits. In the command line mode of scDRS, it also performs extra analysis and reads in files for multiple times, which would further increase its runtime. To isolate the runtime of only the cell type-trait association process, we wrote a new python script that excluded these unrelated steps. All time for reading in the dataset for scDRS was also excluded from the comparison. To control the CPU usage, podman containers were used to limit the CPU usage to a single core.

## Data preprocessing

### GWAS datasets

We processed and analyzed GWAS summary statistics across a total of 30 complex traits (Supplementary Table 1). Because the majority of GWAS summary statistics were reported using the GRCh37 reference build, any studies with summary statistics based on GRCh38 were converted to GRCh37 using LiftOver [51] for consistency. SNP rsID annotations used dbSNP build 151. Duplicated SNPs were dropped and SNPs without annotations in dbSNP were retained and named by their respective chromosomes and positions.

We used MAGMA (v1.09b) to annotate SNPs to genes and compute gene-level z-scores for each trait, which reflect the overall gene-level association to the trait after factors such as linkage disequilibrium and population stratification are regressed out based on individuals of European ancestry from Phase 3 of the 1,000 Genomes Project [52]. SNP assignment to genes used a 35 kb upstream and 10 kb downstream window around the gene body, as recommended in previous studies [6]. We also examined the effects of systematically varying the MAGMA gene window size on *seismic* results (Supplementary Figure 2).

### scRNA-seq datasets

We processed and analyzed 4 large atlas-scale scRNA-seq datasets, specifically Tabula Muris FACS (TM FACS) [14], Tabula Muris droplet (TM droplet) [14], Tabula Sapiens (TS) [19], and Saunders et al. [21], capturing over 1.5 million cells across over 60 tissue regions (Supplementary Table 2). The TM FACS, TM droplet, and TS datasets each had different quality control filters before we downloaded them. Specifically, the two TM datasets had removed mitochondrial genes and filtered outlier cells. We additionally filter out cells with fewer than 2,000 unique molecular identifier (UMI) counts. The TS dataset had filtered out cells with fewer than 2,500 counts but did not consider mitochondrial gene expression, so we filter out cells where >10% counts are from mitochondrial genes. The Saunders et al. dataset provided cell-level annotations for doublets, outliers, singletons, and unannotated cells, all of which we excluded. We further filtered out cells if either >10% of counts were from mitochondrial genes or if there were fewer than 1000 reads in the cell. Raw gene expression counts were normalized using cell-specific size factors estimated by *scran* [53]. Subsequent analyses use log2(normalized counts) with a pseudocount of 1. For all datasets, we filtered out genes that were either expressed in fewer than 10 cells or had lower than 0.01 mean log-expression across all cell types. We also only retained cell types that had at least 20 cells.

For comparisons using mouse scRNA-seq data, we used mouse-to-human gene mappings from biomaRt [50]. Genes without mapping to human genes were discarded, and for genes with multiple mappings to a single human gene, the mean specificity score was used.

We also manually examined the cell type label annotations for all datasets to resolve or filter out cells with unclear annotations. For example, certain cells had clearly confusing labels, such as hepatocytes in the heart tissue in TS and so were excluded from further analyses. Cell types in TM and TS were manually annotated to corresponding terms in the Cell Ontology [54, 55], and unresolvable cell types were excluded.

For the Saunders et al. dataset, cells were annotated to different regions that had been sequenced separately (frontal cortex, posterior cortex, substantia nigra, hippocampus, thalamus, cerebellum, globus pallidus, entopedeuncular nucleus), as well as cell classes, clusters and subclusters. Cell classes represent broad cell types of a region, which Saunders et al. further refined by iterative clustering into progressively more specific cell clusters and subclusters. For neurons, the subclusters had also been more precisely annotated to their respective structure in the region based on not only computational inference, but also immunohistochemical validation. To establish better consistency for cell types at each granularity, we manually cleaned and refined these annotations. Specifically, for the ‘subclass’ granularity, non-neuronal cells retained their corresponding annotations (e.g., microglia, oligodendrocytes), while neuronal cells were categorized by neurotransmitter type (i.e., excitatory, inhibitory, dopaminergic, cholinergic). For the ‘brain region + class’ granularity, cells were grouped by both brain region and broader cell class (i.e., neuronal cells were considered as one entity and not subdivided by neurotransmitter type). Similarly, for the ‘brain region + subclass’ granularity, cells were grouped by both brain region and subclass. For the ‘brain region + cluster’ granularity, cells were grouped by both brain region and cleaned cluster annotations provided by Saunders et al. (which had finer resolution than subclass and corresponds to specific cortical layers or neuronal connectivity). Saunders et al. also further divided clusters into subclusters, the finest resolution annotations that they provide. These subcluster annotations were often redundant, so after resolving typos, we combined subclusters that had closely related annotations when they were derived from the same cluster within a dissected brain region. ‘Brain region + fine cluster’ is the granularity that corresponds to combining these combined subcluster annotations with brain region information. For interneurons, which span multiple brain regions, subclusters were combined by the primary gene marker. The number of cell types thus varied from 14 at the ‘subclass’ granularity to 231 at the ‘brain region + fine cluster’ granularity (Supplementary Table 4).

