## Supplementary Figures for "Disentangling associations between complex traits and cell types with *seismic*"

### Seismic: supplementary materials

**Supplementary Table 1:** Summary information for the GWAS data sets used in Figures 2-5.
See excel file STable 1.

**Supplementary Table 2:** A listing of all single cell RNA-seq expression datasets used in the paper.
See excel file STable 2.

**Supplementary Table 3:** Running time for cell type-trait association methods (*seismic*, scDRS, FUMA, S-MAGMA) as a function of the number of cells used in the single cell RNA-seq input.
See excel file STable 3.

**Supplementary Table 4:** Detailed cell type-trait association results for each of the four methods across 27 GWAS studies using Tabula Muris FACS data.
See excel file STable 4.

**Supplementary Table 5:** Detailed cell type-trait associations for the Parkinson's disease GWAS and the mouse brain scRNA-seq dataset subset at different cell type granularities.
See excel file STable 5.

**Supplementary Table 6:** All results generated for the Alzheimer's case study depicted in Figure 6, including cell type-trait associations, influential gene and GO enrichment results for the selected cell types.
See excel file STable 6.

### Supplementary figures

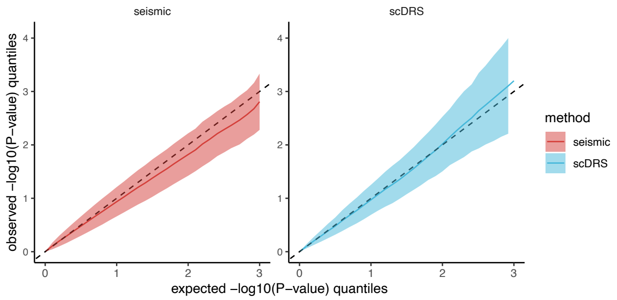


**Supplementary Figure 1:** Quantile plots for the expected P-values versus the observed P-values in null simulation for a random cell type and a random gene sets. In this case the gene index of the MAGMA Z-score vector is shuffled to obtain a random vector to represent the random trait's MAGMA Z-score. The gene index remains constant in the expression matrix. MAGMA software does not accept custom gene Z-score as the input thus only the results of *seismic* and scDRS are shown.

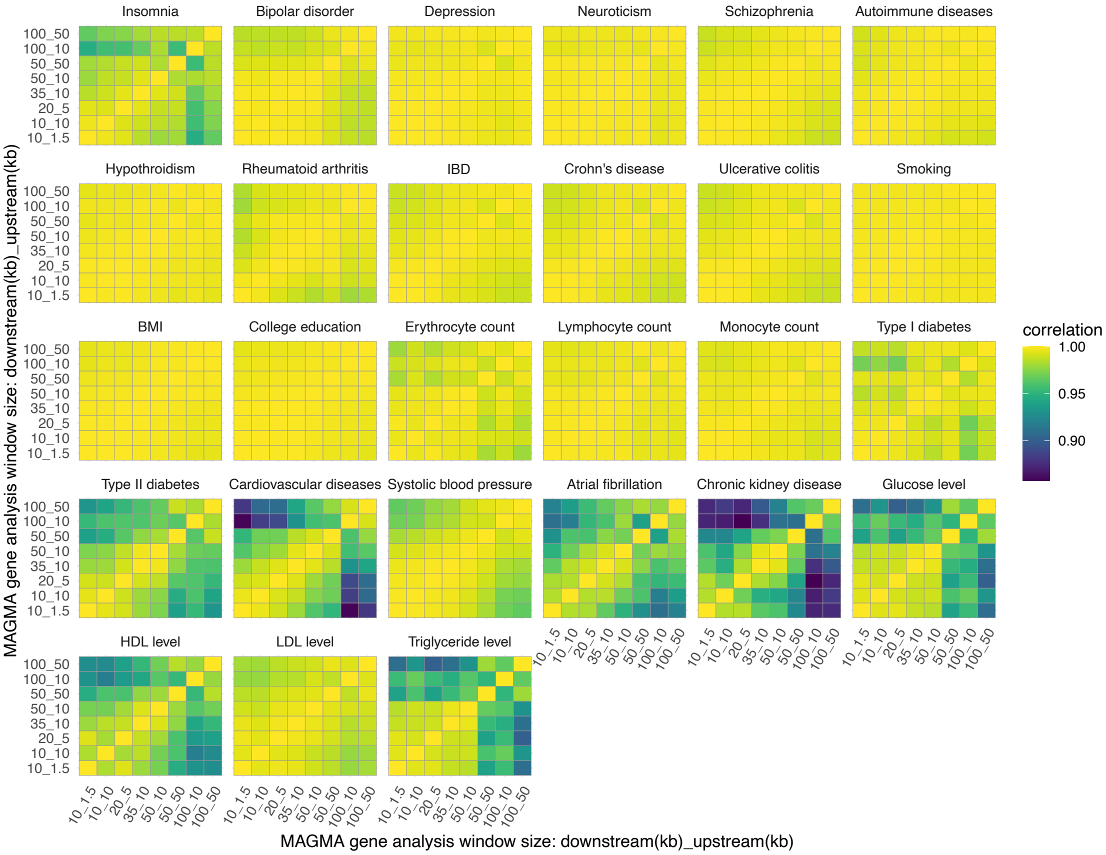


**Supplementary Figure 2:** Correlation of cell type-trait associations across varying MAGMA window sizes. The indices show the range of the gene annotating window (in kilobases). IBD, Inflammatory bowel disease.

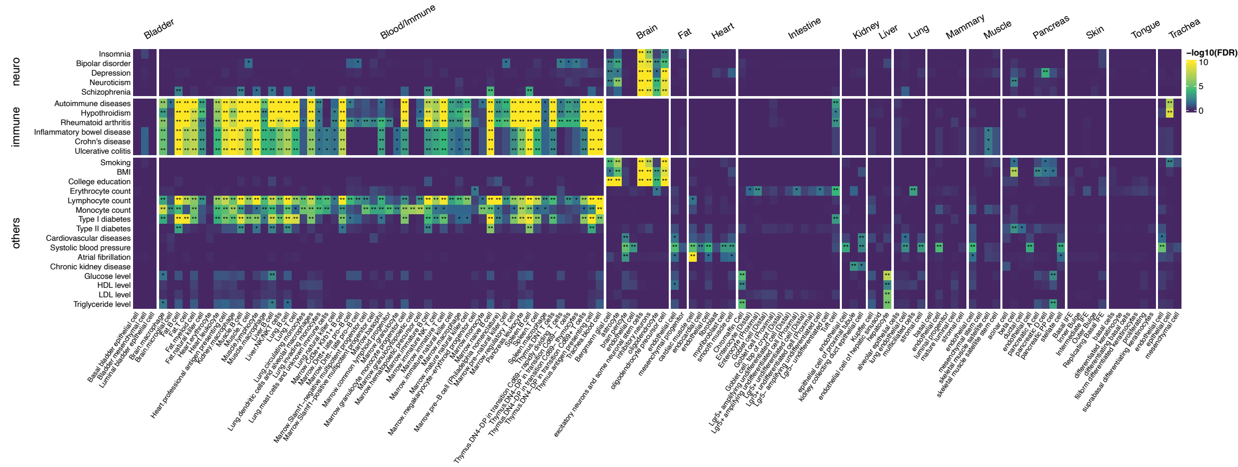


**Supplementary Figure 3:** Heatmap of cell type-trait associations across 27 GWAS identified by scDRS using cell types in Tabula Muris FACS data.

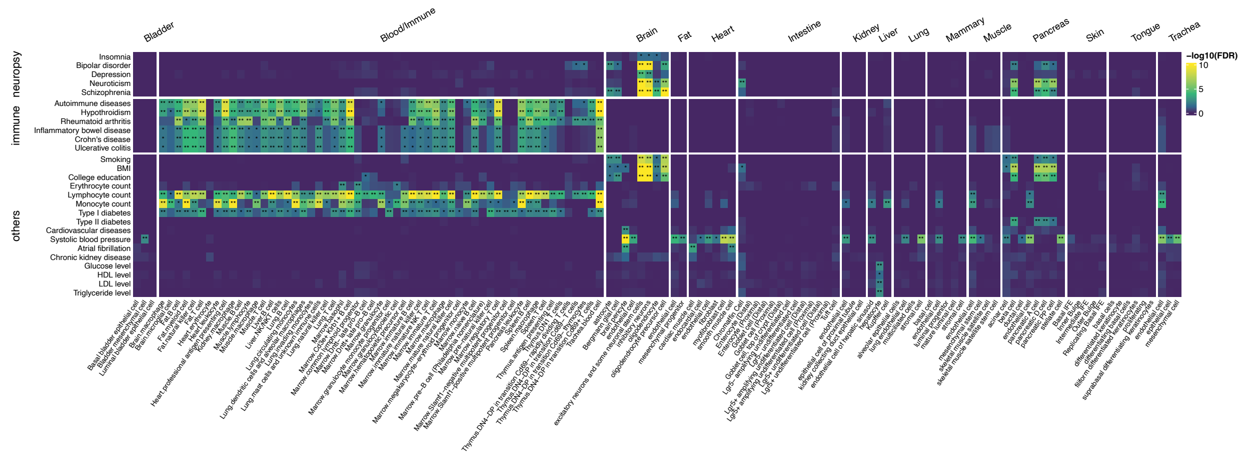


**Supplementary Figure 4:** Heatmap of cell type-trait associations across 27 GWAS identified by FUMA using cell types in Tabula Muris FACS data.

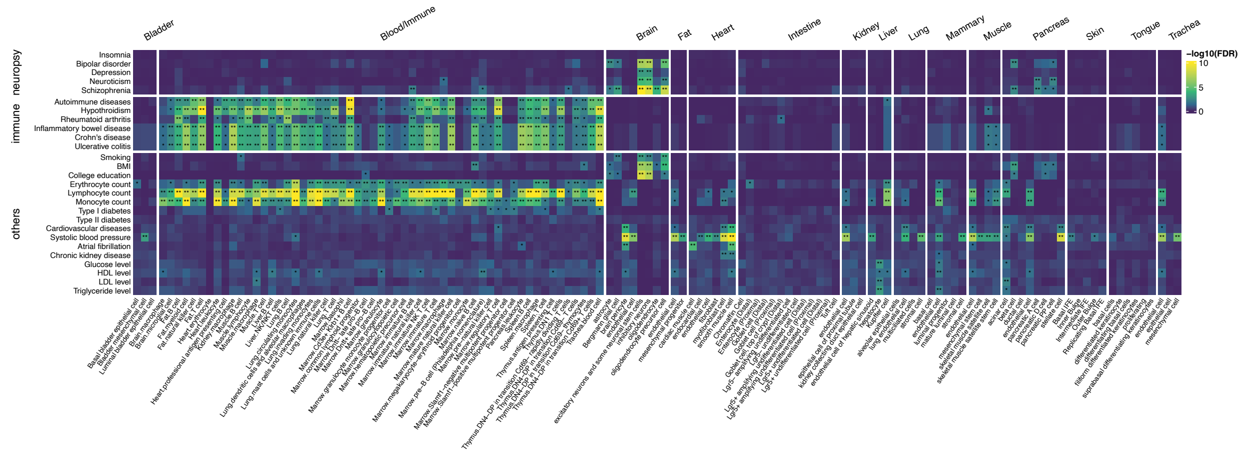


**Supplementary Figure 5:** Heatmap of cell type-trait associations across 27 GWAS identified by S-MAGMA using cell types in Tabula Muris FACS data.

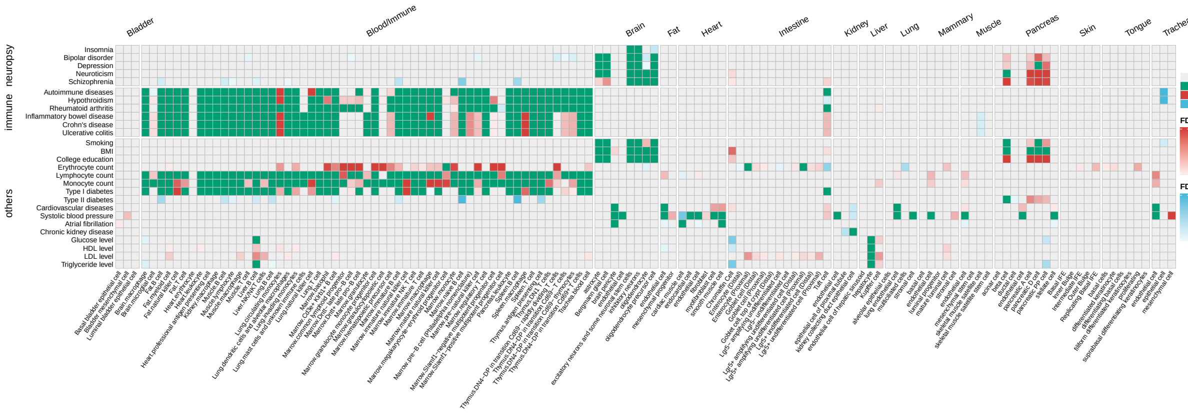


**Supplementary Figure 6:** Heatmaps showing the concordance of results between *seismic* and scDRS across cell type-trait associations pair using the TM FACS data.

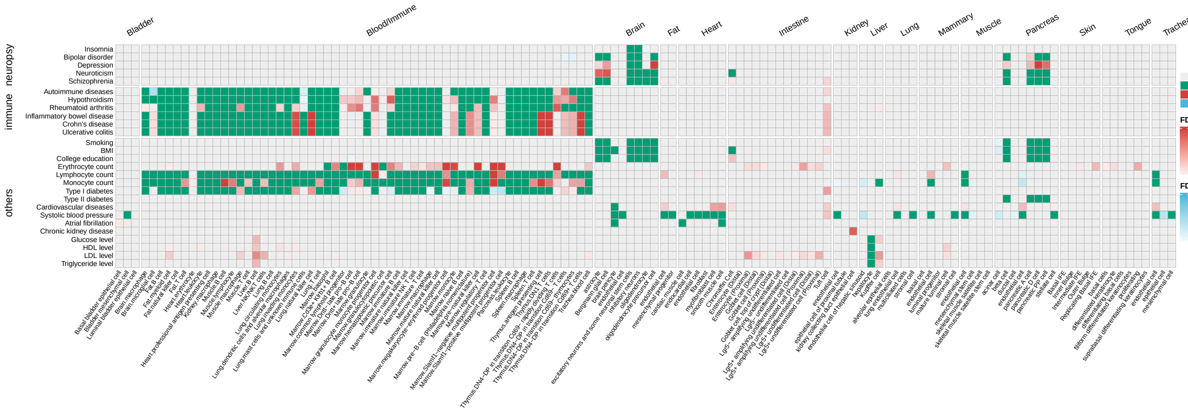


**Supplementary Figure 7:** Heatmaps showing the concordance of results between *seismic* and FUMA across cell type-trait associations pair using the Tabula Muris FACS data.

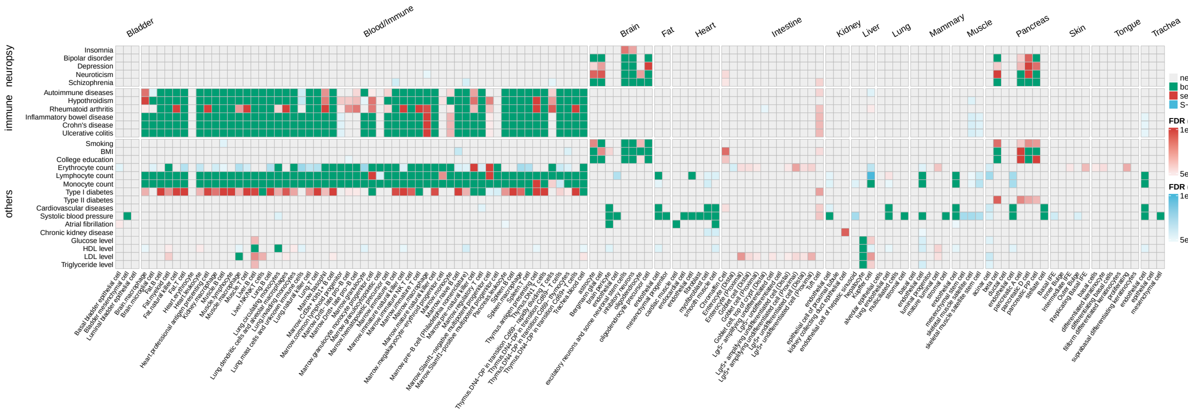


**Supplementary Figure 8:** Heatmaps showing the concordance of results between *seismic* and S-MAGMA across cell type-trait associations pair using the Tabula Muris FACS data.

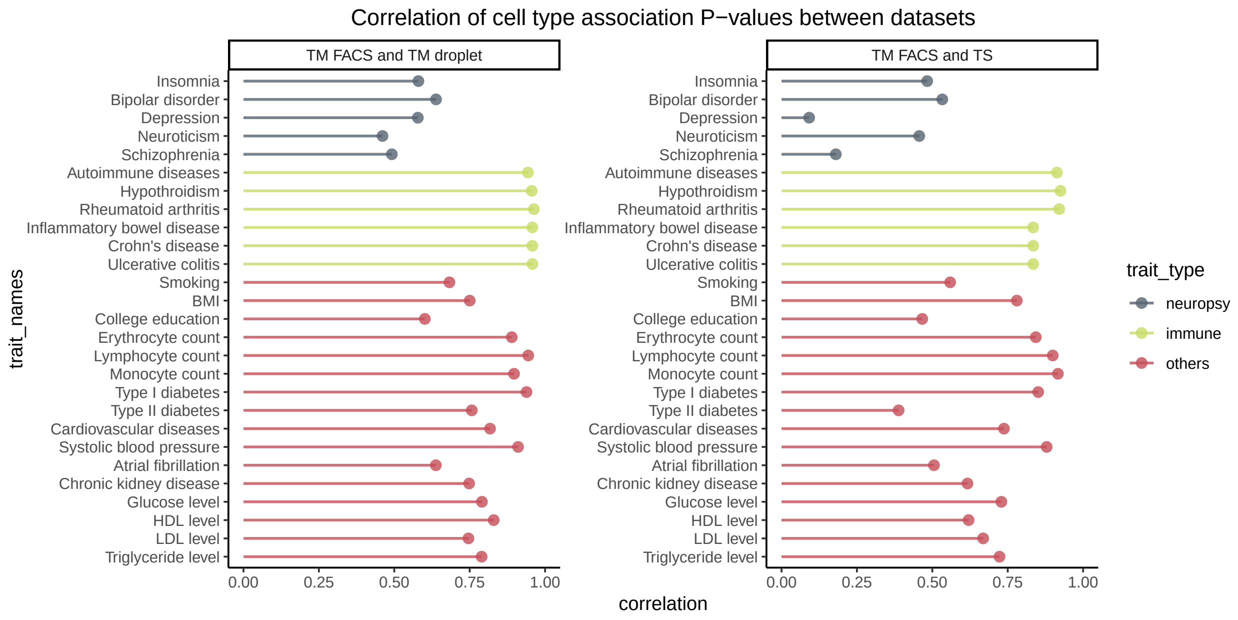


**Supplementary Figure 9:** Spearman's correlations of cell type associations with traits between datasets. The correlation was calculated using only cell types with matching tissue and Cell Ontology IDs.

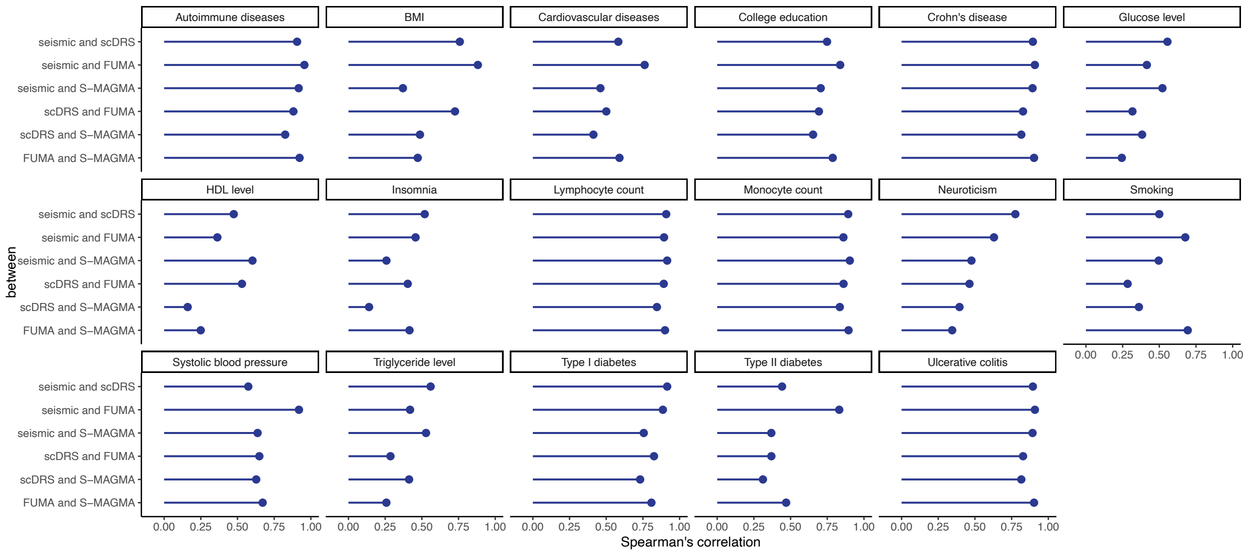


**Supplementary Figure 10:** Pairwise Spearman's correlations for the remaining GWAS traits not shown in Figure 3 using Tabula Muris FACS data.

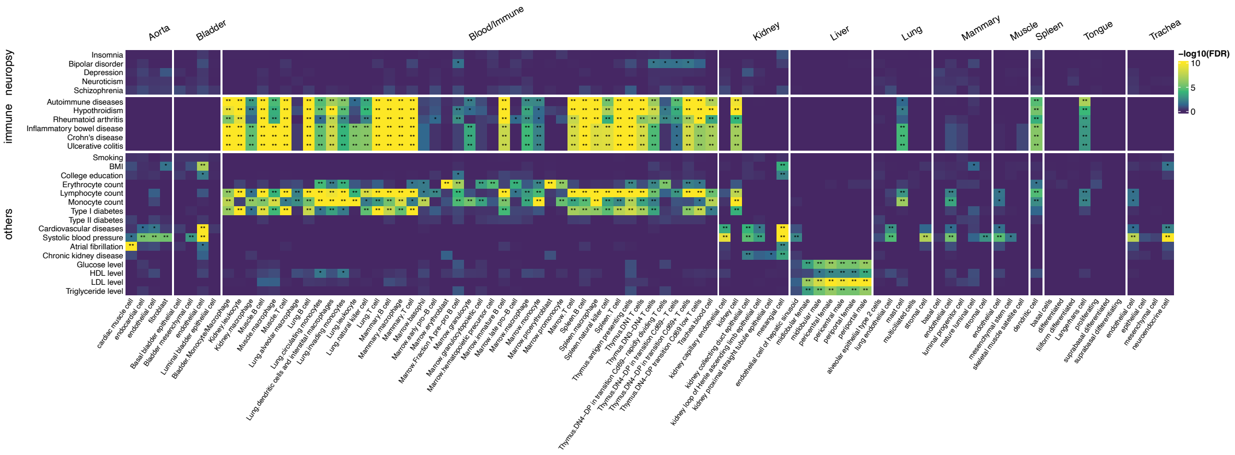


**Supplementary Figure 11:** Heatmap showing cell type-trait associations detected by *seismic* using the Tabula Muris droplet data.

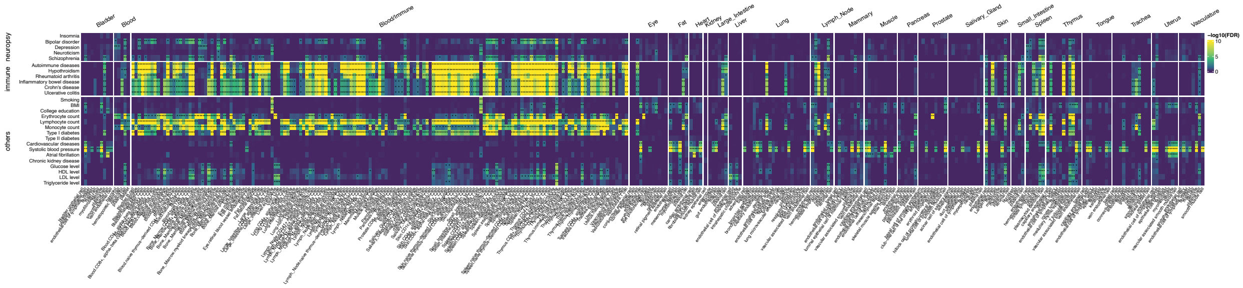

**Supplementary Figure 12:** Heatmap showing cell type-trait associations detected by *seismic* using the Tabula Sapiens dataset.

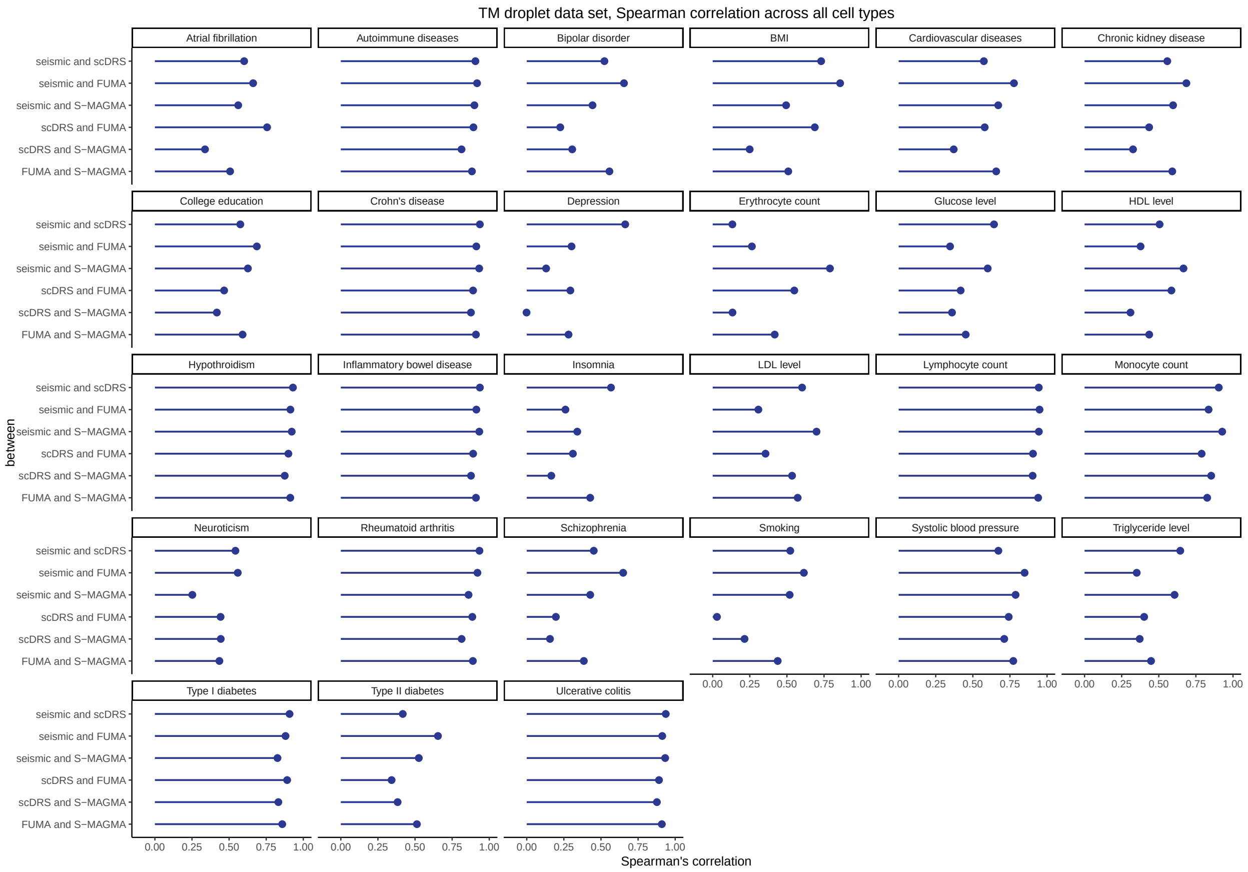


**Supplementary Figure 13:** Pairwise Spearman's correlations for all GWAS traits using Tabula Muris droplet data.

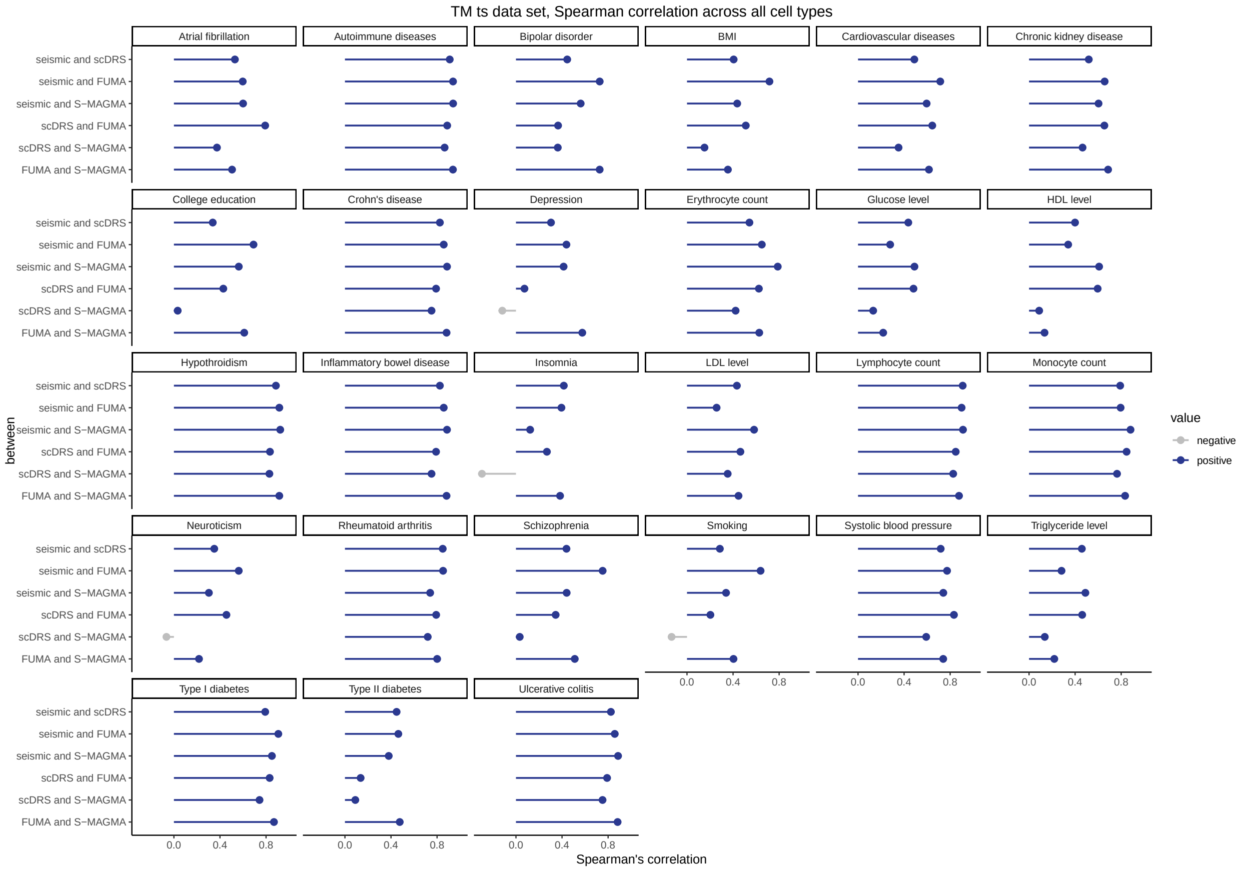


**Supplementary Figure 14:** Pairwise Spearman's correlations for all GWAS traits using the Tabula Sapiens dataset.

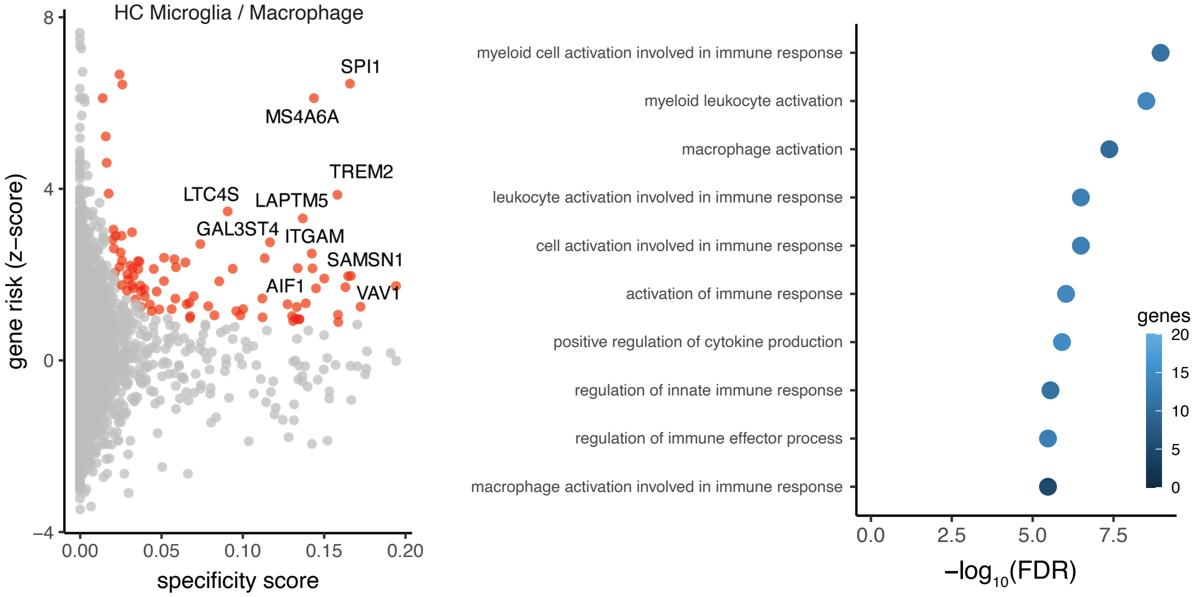


**Supplementary Figure 15:** Influential gene and corresponding GO biological process enrichment for genes driving the association between AD clinical GWAS and the hippocampal microglia / macrophages cell type.
